# Host response against SARS-CoV-2 infection is impacted by allosteric modulation of cholinergic and purinergic receptors

**DOI:** 10.64898/2026.10.22.761280

**Authors:** Victoire Perraud, Guilherme Dias de Melo, Marco Cecchini, Lauriane Kergoat, Julien Lannoy, Jean-Pierre Changeux, Hervé Bourhy, Florence Larrous

**Author notes:** corresponding author *Email address* (F. Larrous).

## Abstract

The COVID-19 pandemic has caused nearly 8 million deaths and a broad spectrum of clinical manifestations driven by dysregulated host inflammation, notably involving key signalling proteins IL-6, CXCL10, and CXCL5. We previously demonstrated that ivermectin (IVM) improves clinical outcomes and reduces the IL-6/IL-10 ratio in the lungs of SARS-CoV-2-infected hamsters. Here, combining high-throughput confocal microscopy and transcriptomics in human cell co-cultures and a hamster model, we dissected how IVM modulates host cells responses via α7nAChR and purinergic receptors. *In vitro*, IVM acted independently of α7nAChR, instead triggering a P2X4-dependent drop in intracellular ATP, rapidly upregulating CXCL10 in macrophages, and suppressing CXCL5 and IL-6. *In vivo*, however, α7nAChR antagonists abolished IVM’s clinical benefit, as did the P2X4 blocker BX430; moreover, whereas the orthosteric agonist EVP-6124 was ineffective, the positive allosteric modulators (PAMs) IVM and GAT-107 reduced lung pathology and anosmia. Thus, IVM operates through a dynamic, context-dependent interplay: P2X4 drives early innate responses in cellular systems including macrophages, whereas the situation *in vivo* is more complex with α7nAChR being essential for tissue protection. These data position selective PAMs and the CXCL10/CXCL5 axis as promising targets for host-directed therapy in COVID-19.

**Author summary:** COVID-19 has killed millions of people worldwide, and much of its harm comes not from the virus itself but from the patient’s own immune response spiralling out of control. To understand how this inflammation is controlled, we used ivermectin (a well-known anti-parasitic drug) as a research tool, building on our earlier finding that it improved the health of SARS-CoV-2-infected hamsters. In human immune cells grown in the laboratory, ivermectin modulated an ATP-sensing receptor, protected these cells, and shifted the inflammatory signals they released. In infected hamsters, however, protecting the animals’ health and their sense of smell required both this receptor and a second one, which responds to acetylcholine, one of the body’s natural chemical messengers, while protecting the lungs relied in particular on this second receptor. These findings do not support treating patients with ivermectin. Rather, they map the machinery behind harmful inflammation and point to a promising path forward: purpose-built drugs that fine-tune the body’s immune response in COVID-19 and other inflammatory diseases.

## Introduction

The COVID-19 pandemic, which emerged in December 2019, caused nearly 800 million cases worldwide, including almost 8 million deaths, since its beginning (1). Once infected, patients can present different clinical profiles, ranging from asymptomatic to severe and even fatal forms. The most common symptoms are not specific to the disease and overlap with those of the flu-like syndrome (fever, sore throat, cough, headache, myalgia and asthenia) (2). Some patients with mild symptoms may also experience neurological symptoms such as dizziness, ageusia and anosmia (3–5). During SARS-CoV-2 infection, an up-regulation of multiple pro-inflammatory cytokines (IL-6, CXCL10, CXCL1, CXCL2 and CXCL5) was observed (6), in which CXCL10 is involved in monocyte stimulation and macrophage chemoattraction (7), and CXCL5 in neutrophil chemotaxis (8).

Although several approved drugs and experimental products have shown antiviral activity against SARS-CoV-2 *in vitro*, at present there are only few antiviral therapies that have been authorized or approved to treat mild to moderate COVID-19 in people who are more likely to get very sick. These therapies include agents issued from drug repurposing such as inhibitors of RNA virus replication and inhibitors of HIV replication, as well as more specific developments, such as potent inhibitor of SARS-CoV 3C-like protease and a nucleoside analogue prodrug that shows inhibitory effects on pathogenic animal and human coronaviruses (9, 10)

Ivermectin (IVM), an anthelmintic drug, has immuno0modulating properties in animals and humans (11). Our group and others have shown that during SARS-CoV-2 infection in animal models, IVM plays an immunomodulatory role, where the IL6/IL10 ratio was considerably reduced in lung tissue, probably explaining the more favourable clinical presentation of treated animals (12). Furthermore, prophylactic treatment with IVM in a hamster model resulted in a reduction in clinical signs and inflammation (13). Although the exact mechanisms of IVM action are still unknown (14), it may involve the α7 nicotinic acetylcholine receptor (α7nAChR), known to mediate anti-inflammatory responses through a reduction of pro-inflammatory cytokines, chemokines and activation of immune cells (15).

IVM is a member of the type I positive allosteric modulators (PAM) of α7nAChR (16, 17). Type I PAMs bind to the allosteric site of the receptor, distinct from the orthosteric site, which is recognized by the endogenous agonist. Type I PAMs potentiate the receptor response when applied at the same time as agonists, but do not slow down desensitization, contrary to type II PAMs (18).

Additionally, IVM is also a positive allosteric modulator of purinergic receptors P2X4 and P2X7 receptors. IVM facilitates the opening and retards the closing of the channel of both P2X4 (19) and P2X7 receptors (human, but not rat or mouse) (20). These receptors are activated by extracellular ATP (21, 22), which can be released by dying or lysed cells. In response to the ATP signal, purinergic receptors can consolidate the inflammatory response (Di Virgilio et al., 2020) and could play a role during SARS-CoV-2 infection. Indeed, to control the inflammation induced by SARS-CoV-2, drugs with inhibitory effects on purinergic receptors were proposed, such as lidocaine (23), an antagonist of P2X7 receptor, or fluoxetine and paroxetine, antidepressants which are P2X4 inhibitors (24). Moreover, P2X4 and P2X7 are co-expressed on epithelial, immune and inflammatory cells (25) and exhibit functional interactions (26–28). However, the data regarding the functional interaction between P2X4 and P2X7 subunits are contradictory. Currently, despite efforts to determine whether P2X4 and P2X7 interact as heterotrimers or homotrimers, there is no definitive evidence in favour of either model (29).

Although IVM has never been formally recommended in humans in prevention or treatment of COVID-19 based on the various clinical trials conducted (30–32), preclinical findings in hamsters warrant further investigation to understand its mechanism of action. Moreover, it was shown that α7nAChR, P2X4 or P2X7 receptors can collaborate in rats or mice, through interaction or through Panx1 channels (33–35). Those receptors could also be involved in the pathogenesis of SARS-CoV-2 infection (36).

In this study, we sought to investigate the role of α7nAChR and P2X4 in IVM’s modulation of the host response during SARS-CoV-2 infection. By using high-throughput confocal microscopy, complex co-culture systems and transcriptomic analysis, it appears that IVM operates independently to α7nAChR through a dynamic, context-dependent interplay where P2X4 drives early innate responses in cellular systems. However, the situation *in vivo* is more complex with α7nAChR being essential for tissue protection. Finally, our results indicate that IVM’s immunomodulation is mostly achieved through CXCL10 and CXCL5 pathways and involves a dynamic interplay among multiple cell types.

## Results

### Modulation of α7nAChRs is not sufficient to regulate SARS-CoV-2 infection *in vitro*

We have previously shown *in vivo* that IVM exerts an immunomodulatory role leading to improved clinical outcomes in treated animals (12). Based on the nicotinic hypothesis (37) and the established role of nicotinic receptors in the modulation of the cholinergic anti-inflammatory pathway (38), we sought to determine whether α7nAChR contributes to the modulation of host response during SARS-CoV-2 infection in a cellular model. To specifically investigate the role of α7nAChRs activation, we selected two additional modulators of the α7nAChRs from the Nicotinic Acetylcholine Receptor Allosteric Ligand Library (ACRALL) (39, 40): a PAM capable to independently activate the receptors on its own (ago-PAM), GAT-107 (GAT) (41) and a competitive agonist, EVP-6124 (EVP) (42).

We established a simplified *in vitro* model in which Vero E6 cells were infected with the ancestral SARS-CoV-2 Wuhan. IVM treatment (5 µM) significantly reduced cell death and the percentage of infected cells compared to untreated controls (Figure 1A and 1D; Figure S1A–B). Similar protective effects were observed with other viral variants, including Alpha, Beta, and Delta (Figure S1C–H), indicating that the underlying mechanism is variant-independent. We next tested the other selected α7nAChR modulators against SARS-CoV-2 Wuhan. GAT (25 µM) slightly increased cell viability (Figure 1A; Figure S1I) but significantly reduced the number of infected cells (Figure 1D; Figure S1J). Likewise, EVP (16 µM) significantly increased total cell counts (Figure 1A; Figure S1K) and decreased the proportion of infected cells (Figure 1D; Figure S1L).

**Figure 1:**
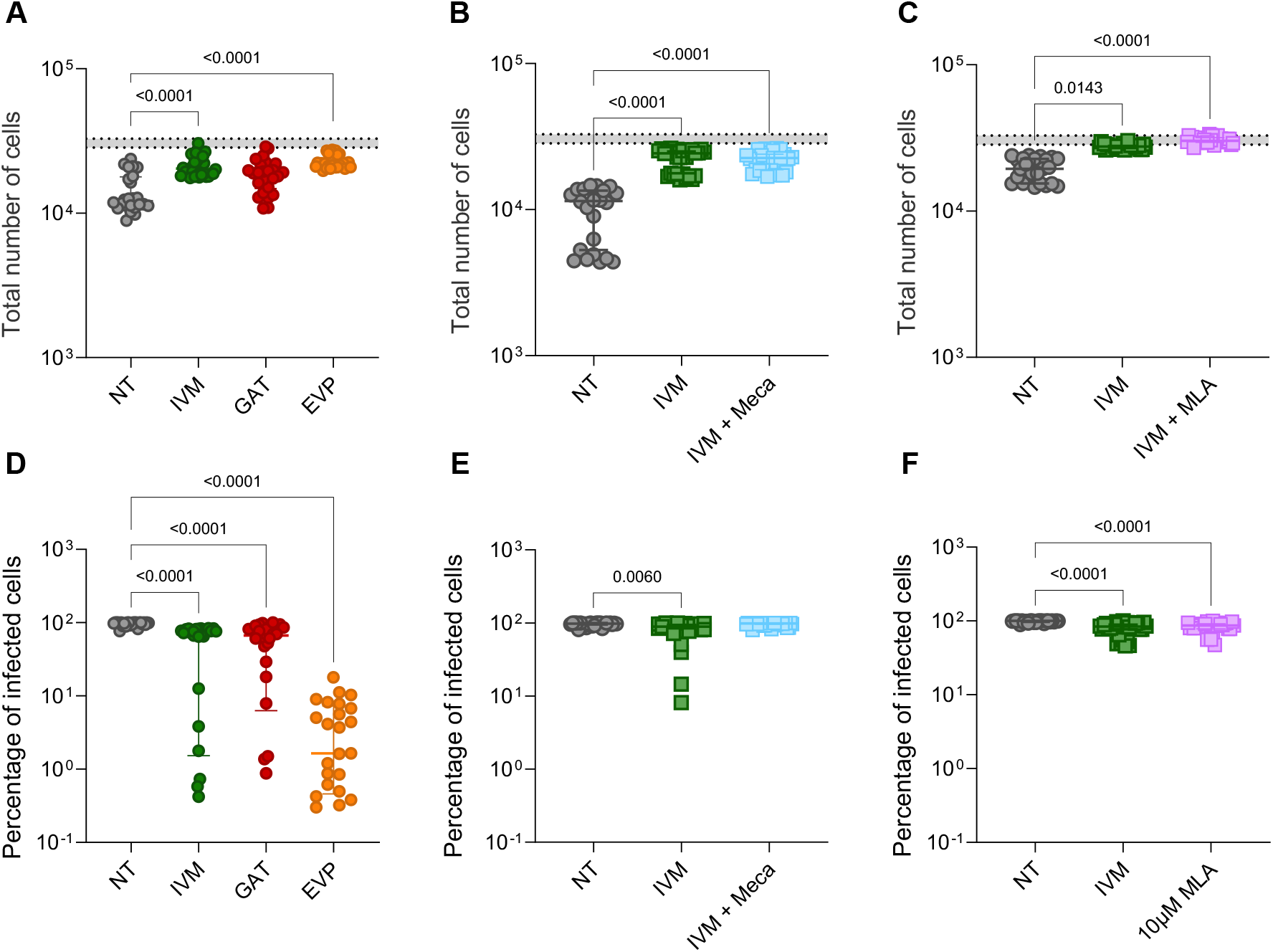
Modulation of α7nAChRs is not sufficient to regulate SARS-CoV-2 infection *in vitro*. Effects of α7nAChR modulators or antagonists on monocellular *in vitro* model using Vero E6 cells and the Opera Phenix High Content Screening System technology (only the infected conditions are shown). **(A)** Total number of Vero E6 cells and **(D)** percentage of infected Vero E6 cells present after 56 h of infection (MOI 0.1) with the original virus (Wuhan) treated with 5 µM of ivermectin (IVM), 25 µM of GAT and 16 µM of EVP. **(B, C)** Total number of cells treated with ivermectin plus **(B)** 32 µM Meca or **(C)** 10 µM MLA. **(E, F)** Percentage of infected cells treated with ivermectin plus **(E)** 32 µM Meca or **(F)** 10 µM MLA. All the values are expressed as median, error bars indicate ± interquartile range. Kruskal-Wallis test followed by the Dunn’s multiple comparisons test (only the significant values are shown). Three independent experiments with 10 technical replicates were performed with the Opera Phenix technology. Four independent experiments in triplicates were performed for the Nluc and ATP luminescence, where the data are normalized on the infected non-treated condition.

Since IVM, GAT, and EVP all conferred a relative cellular protection in this model, we next assessed whether this effect was dependent on α7nAChR activation. We therefore combined IVM with two structurally distinct nAChR antagonists with the aim to abrogate the protective effect of IVM: mecamylamine (Meca), a non-competitive non-specific antagonist, and methyllycaconitine (MLA), a competitive antagonist that is highly selective for α7nAChR. However, neither MLA (0.1–10 µM) nor Meca (8–32 µM) abolished the protective effect of IVM (Figure 1B, 1C, 1E, 1F; Figure S2).

Taken together, these results show that, although activation of α7nAChR can protect cells from SARS-CoV-2 infection, IVM exerts its protective effect via an α7nAChR-independent mechanism, pointing to a distinct, probably multimodal, mode of action.

### IVM exerts a protective effect against SARS-CoV-2 infection *in vitro* via an α7nAChR-independent mechanism

To further characterize the effects of α7nAChR modulators, Vero E6 cells were infected with a recombinant SARS-CoV-2 expressing nanoluciferase (Wuhan-nLuc) and treated. At 56 h post-infection (hpi), intracellular ATP levels and nanoluciferase activity in cell supernatants were measured. Intracellular ATP was approximately 3-fold higher in non-infected than in infected cells, validating it as a robust indicator of cell integrity. Nanoluciferase release in the supernatant correlated with the level of infection. Upon treatment with IVM, GAT, or EVP, intracellular ATP was higher (Figure 2A) and nanoluciferase luminescence was significantly reduced compared with untreated infected controls (Figure 2B), confirming the protective effects observed previously.

**Figure 2:**
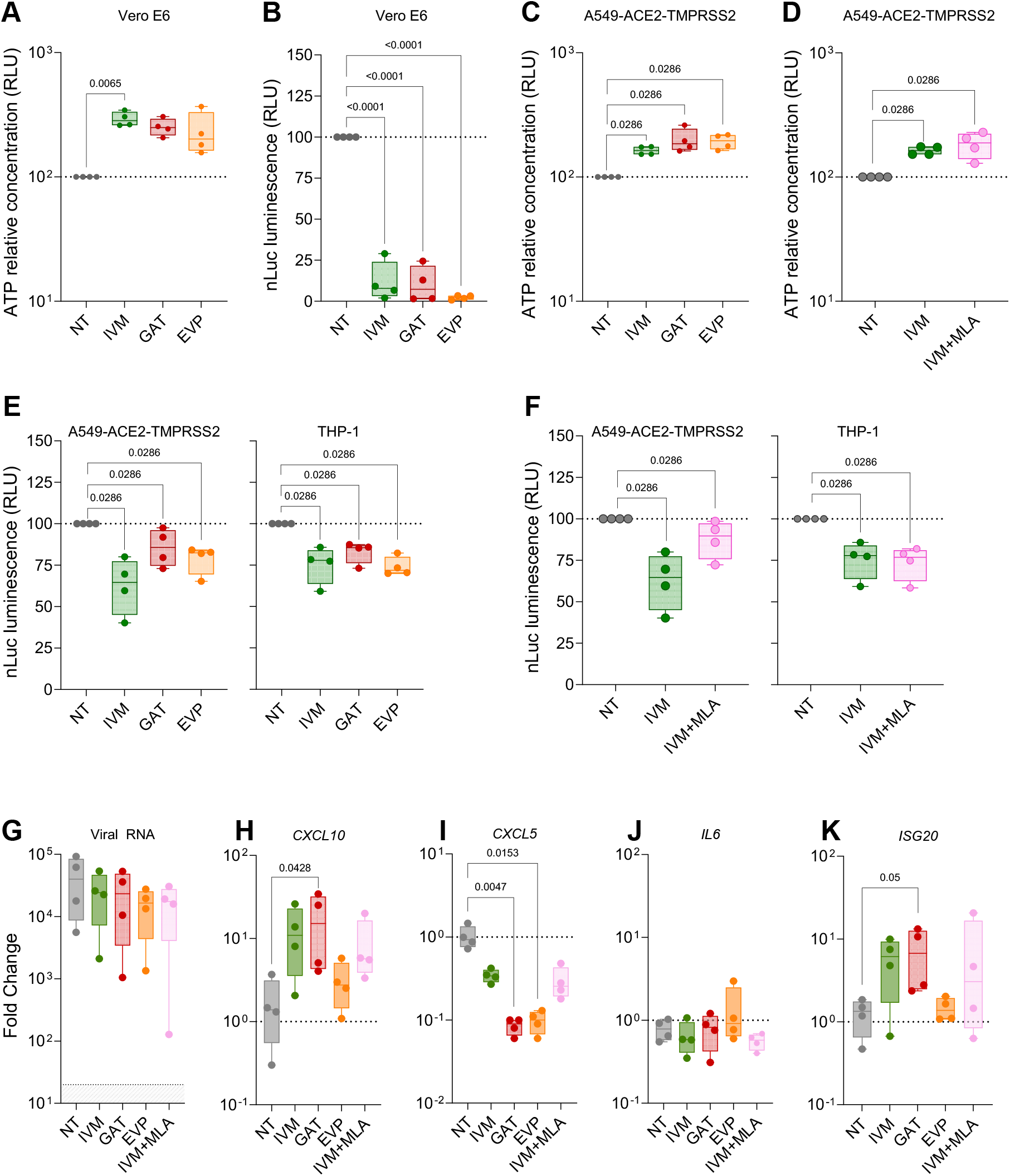
IVM exerts a protective effect against SARS-CoV-2 infection *in vitro* via an α7nAChR-independent mechanism. In a single cell model, Vero E6 cells were infected with SARS-CoV-2/Wuhan_nLuc (MOI 0.1) for 56 hpi. **(A)** ATP relative concentration measured in Vero-E6 cells. **(B)** Nanoluciferase (Nluc) luminescence resulting from Wuhan_nLuc virus (MOI 0.1) quantified in cells supernatant. In the Transwell model, A549-ACE2-TMPRSS2 cells were infected at an MOI of 0.1 with a Wuhan-nLuc virus and co-cultured with THP-1 cells. **(C)** ATP relative concentration of infected A549-ACE2-TMPRSS2 cells measured at 72 hpi in the presence of α7nAChR modulators (5 µM IVM, 25 µM GAT or 16 µM EVP) or α7nAChR antagonists (10 µM MLA) **(D)**. Nanoluciferase (Nluc) luminescence of infected A549-ACE2-TMPRSS2 and THP-1 cells measured at 72 hpi in the presence of α7nAChR modulators (5 µM IVM, 25 µM GAT or 16 µM EVP) **(E)** or α7nAChR antagonist (10 µM MLA), **(F).** The values are normalized on infected non-treated conditions, only the infected conditions are shown. Four independent experiments were performed in technical triplicate. All the values are expressed as median, error bars indicate ± interquartile range. Mann-Whitney test was performed (P<0.05). The expressions of **(G)** viral RNA and **(H)** *CXCL10*, **(I)** *CXCL5,* **(J)** *IL6*, and **(K)** *ISG20* mRNA were measured in THP-1 cells at 72 hpi. The values are normalized on non-infected non-treated conditions, only the infected conditions are shown. Four independent experiments were performed in technical duplicates. All the values are expressed as median, error bars indicate ± interquartile range. Mann-Whitney test was performed (P<0.05). The statistical test indicated with p value is performed between the non-treated condition and the different treated ones.

To gain further insight into the mechanism of action of IVM, we developed a co-culture model combining epithelial and macrophage cell lines. A549 cells, a type of human alveolar basal epithelial cell, stably expressing ACE2 and TMPRSS2 (A549-ACE2-TMPRSS2) were infected and co-cultured with differentiated THP-1 cells, a human monocyte cell line, in a transwell system, then exposed to IVM, GAT, and EVP for 72 hours. To determine whether the protective effect of IVM was exclusively mediated by α7nAChR, we combined it with MLA, a highly selective competitive antagonist of this receptor.

As before, intracellular ATP was measured to assess cell viability (Figure 2C), while infection was monitored in each compartment using the Wuhan_nLuc reporter (Figure 2E–F). Cholinergic modulators significantly increased intracellular ATP (Figure 2C), reflecting improved cell survival, and this effect was not abolished by co-treatment with MLA (Figure 2D). Similarly, nanoluciferase release was significantly reduced in both A549-ACE2-TMPRSS2 and THP-1 cells (Figure 2E), indicating limited viral replication. Remarkably, MLA partially inhibited the effect of IVM in A549-ACE2-TMPRSS2 cells, but not in THP-1 cells (Figure 2F).

Furthermore, we observed similar viral RNA load in THP-1 cells across all conditions, irrespective of treatment. Notably, α7nAChR modulators induced a distinct transcriptomic signature in THP-1 cells, characterized by increased CXCL10 and decreased CXCL5 gene expression after 72 hpi (Figure 2H–I). Surprisingly, the expression of IL-6 was not affected (Figure 2J), whereas ISG20 was upregulated (Figure 2K). This transcriptional profile remained unchanged when MLA was co-administered with IVM (Figure 2G–K).

Collectively, these results indicate that although IVM exerts a protective effect against SARS-CoV-2, its mechanism of action is not exclusively dependent on α7nAChR. The cell-type-specific sensitivity to MLA further suggests that IVM may act through distinct, and possibly multiple, pathways in epithelial and immune cells.

### Purinergic receptors are also involved in cellular integrity during SARS-CoV-2 infection

Given that IVM is also a positive allosteric modulator of purinergic receptors (19, 20), we hypothesized that P2X4 and/or P2X7 might contribute to its protective effects during SARS-CoV-2 infection. We therefore used selective antagonists of purinergic receptors, BX430, a specific P2X4 antagonist, and A438079, a specific P2X7 antagonist, to evaluate the contributions of these receptors to cell integrity and host immune response (Figure 3A–G).

**Figure 3:**
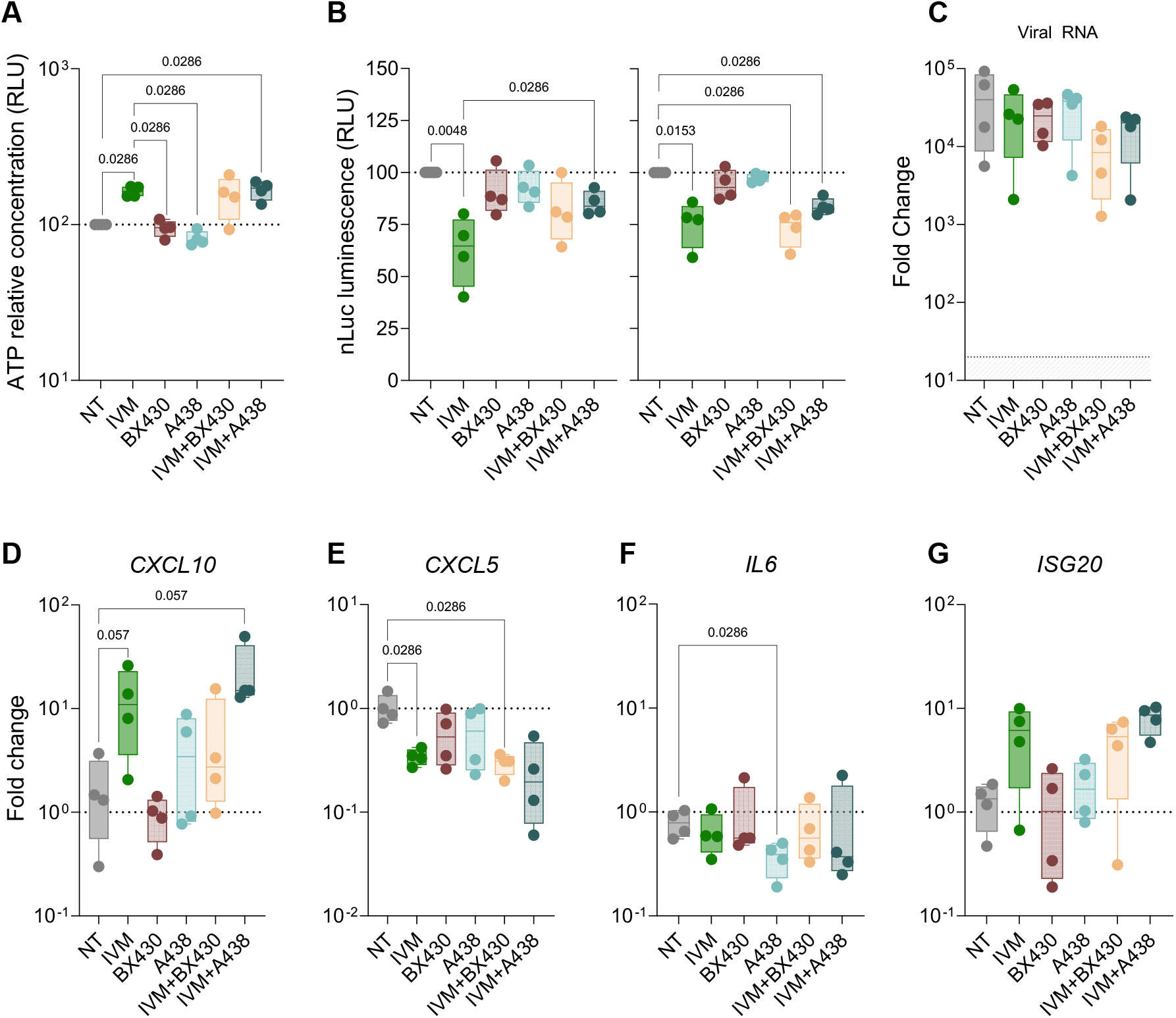
Purinergic receptors are also involved in cellular integrity during SARS-CoV-2 infection. In the Transwell model, A549-ACE2-TMPRSS2 cells were infected at an MOI of 0.1 with a Wuhan-nLuc virus and co-cultured with THP-1 cells, in the presence of ivermectin (IVM) in combination or not with purinergic antagonists: BX430 (at 5 µM, antagonizing P2X4), A438097 (A438, at 10 µM, antagonizing P2X7). **(A)** ATP relative concentration of infected A549-ACE2-TMPRSS2 cells measured at 72 hpi. **(B)** Nanoluciferase (Nluc) luminescence of infected A549-ACE2-TMPRSS2 (on the left) and THP-1 cells (on the right) measured at 72 hpi. Transcriptomic analysis of THP-1 and A549-ACE2-TMPRSS2 cells after 72 hpi with ivermectin (IVM) in combination or not with purinergic antagonists: BX430 (at 5 µM, antagonizing P2X4), A438097 (A438, at 10 µM, antagonizing P2X7). The expression of **(C)** viral RNA and **(D)** *CXCL10*, **(E)** *CXCL5,* **(F)** *IL6*, and **(G)** *ISG20* mRNA expression were measured. The values are normalized on non-infected non-treated conditions, only the infected conditions are shown. Four independent experiments were performed in technical duplicates. All the values are expressed as median, error bars indicate ± interquartile range. Mann-Whitney test was performed (P<0.05). The statistical test indicated with p value is performed between the non-treated condition and the different treated ones.

Treatment with BX430 or A438079 alone had no effect on intracellular ATP levels in A549-ACE2-TMPRSS2 cells compared with untreated infected controls (Figure 3A). Similarly, nanoluciferase release from A549-ACE2-TMPRSS2 or THP-1 cells remained comparable to that of untreated infected cells (Figure 3B), unlike the reduction observed with IVM alone. When combined with IVM, neither BX430 nor A438079 abolished the protective effect on cell viability, as reflected by unchanged intracellular ATP levels in infected A549-ACE2-TMPRSS2 cells (Figure 3A). However, co-treatment with IVM and BX430 or A438079 partially reversed the antiviral effect of IVM in epithelial cells, resulting in increased nanoluciferase release compared with IVM alone (Figure 3B), reminiscent of the partial reversal observed with MLA (Figure 2F). Notably, whereas MLA acted only on A549-ACE2-TMPRSS2 cells, purinergic antagonists acted on both A549-ACE2-TMPRSS2 and THP-1 cells (Figure 3B).

Regarding the transcriptomic profile of THP-1 cells at 72 hpi, neither BX430 nor A438079 alone altered gene expression compared with untreated controls. Surprisingly, the characteristic transcriptomic signature induced by IVM was maintained even in the presence of BX430 or A438079 (Figure 3D–G).

Collectively, these data indicate that both purinergic and cholinergic receptors contribute to the protective effect of IVM during SARS-CoV-2 infection *in vitro*. However, the persistence of the IVM-induced transcriptomic signature despite the presence of purinergic antagonists suggests that the immunomodulatory effects of IVM may involve distinct, and possibly parallel, pathways.

### IVM-mediated immunomodulation reveals distinct contributions of α7nAChR and purinergic receptors

To determine how IVM modulates the host response during SARS-CoV-2 infection and to identify the underlying receptor-dependent pathways, we analyzed the A549-ACE2-TMPRSS2 and THP-1 co-cultures during early time-points post-infection, at 24 and 48 hpi. Viral RNA levels were comparable across all conditions, including IVM alone and IVM combined with nAChR or purinergic antagonists (Figure 4A). However, the transcriptomic analysis revealed a marked temporal and cell-specific dichotomy. THP-1 cells responded to IVM treatment, with significant up regulation of *CXCL10* and downregulation of *CXCL5* and *IL-6* as early as 24 hpi (p=0.0286 for each, Mann-Whitney test) (Figure 4B–D). In contrast, A549-ACE2-TMPRSS2 cells exhibited no significant modulation of *CXCL5* or *IL-6* at either time point and a delayed transcriptomic change, with significant *CXCL10* induction only at 48 hpi (Figure 4B–D). IVM did not induce any significant change in the regulation of *ISG20*, but an upward trend is visible at 48 hpi in A549-ACE2-TMPRRS2 and THP-1 cells.

**Figure 4:**
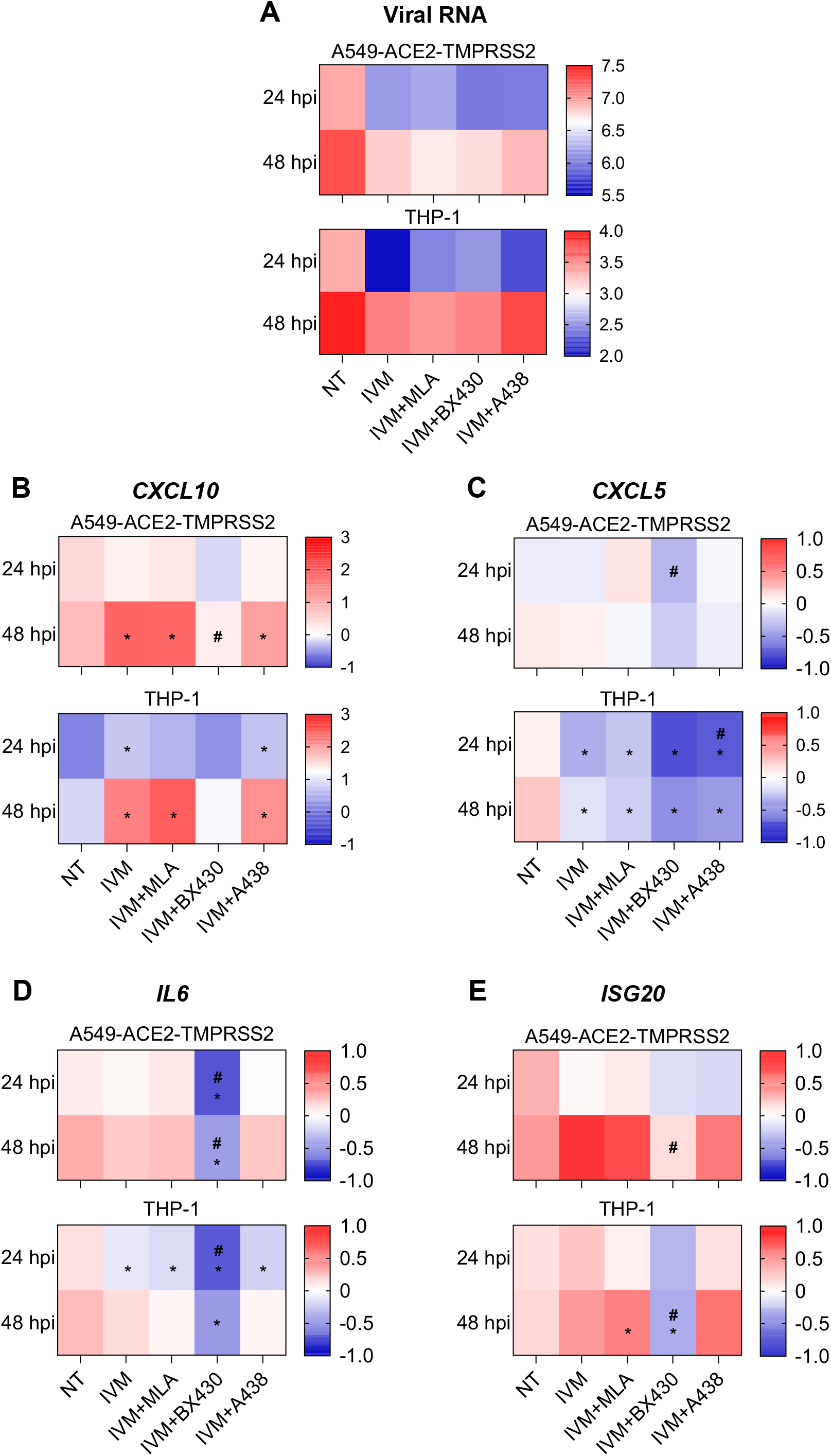
Mechanistic dissection of IVM-mediated immunomodulation reveals distinct contributions of α7nAChR and purinergic receptors. In the Transwell model, A549-ACE2-TMPRSS2 cells were infected at an MOI of 0.1 with a Wuhan-nLuc virus and co-cultured with THP-1 cells, in the presence of ivermectin (IVM) in combination or not with cholinergic or purinergic antagonists: MLA (at 10 µM, antagonizing α7nAChR), BX430 (at 5 µM, antagonizing P2X4), A438097 (A438, at 10 µM, antagonizing P2X7). Transcriptomic analysis was performed on A549-ACE2-TMPRSS2 and THP-1 cells. The expressions of **(A)** viral RNA and **(B)** *CXCL10*, **(C)** *CXCL5,* **(D)** *IL6*, and **(E)** *ISG20* mRNA were measured. The values are normalized on non-infected non-treated conditions, only the infected conditions are shown as heat Map. Four independent experiments were performed in technical duplicates. Mann-Whitney test was performed (P<0.05). The statistical test indicated with p value is performed between the non-treated condition and the different treated ones (*) or between IVM and IVM + antagonist (#).

Blocking α7nAChR with MLA did not alter this IVM-induced transcriptomic signature (Figure 4B–D), demonstrating that the IVM-induced transcriptional immunomodulation is α7nAChR-independent. In contrast, the P2X4 antagonist BX430 prevented IVM-induced *CXCL10* upregulation in both cell types at 48 hpi (Figure 4B). BX430 also further significantly reduced *IL-6* transcription in THP-1 cells at 24 hpi (p=0.0286, Mann-Whitney test) and *IL-6*, and *ISG20* transcription in A549-ACE2-TMPRSS2 cells at 48 hpi (p=0.0286, Mann-Whitney test) (Figure 4B-E). Finally, the P2X7 antagonist A438079 did not modify IVM effects on both types of cells except for *CXCL5* which is significantly further decreased in THP-1 cells beyond the level achieved by IVM alone (p=0.0286, Mann-Whitney test), suggesting a specific, additive role for P2X7 in *CXCL5* regulation (Figure 4C).

Collectively, these results indicate that IVM elicits a rapid and complex macrophage immunomodulatory response rather than α7nAChR, with a discrete contribution of P2X7 to *CXCL5* modulation and a delayed epithelial response mediated through P2X4.

### α7nAChR and P2X4 receptors mediate IVM-induced protection against SARS-CoV-2 *in vivo*

We next sought to determine the potential respective contribution of α7nAChR and P2X4 receptors to the protective effects of IVM *in vivo* using the golden hamster model. As expected, and already shown (12), IVM markedly reduced clinical scores, anosmia incidence and lung pathology in infected animals (Figure 5A-C and S3). Treatment with GAT-107, an ago-PAM of α7nAChR, also improved these outcomes, reducing the clinical score and the proportion of animals with olfactory dysfunction (11.1% versus 68.8% in non-treated animals) (Figure S3), although less potently than IVM (37.5% dysfunction; Figure 5B). In contrast, treatment with EVP-6124, an agonist of α7nAChR showed no effect *in vivo* (Figure S3).

**Figure 5:**
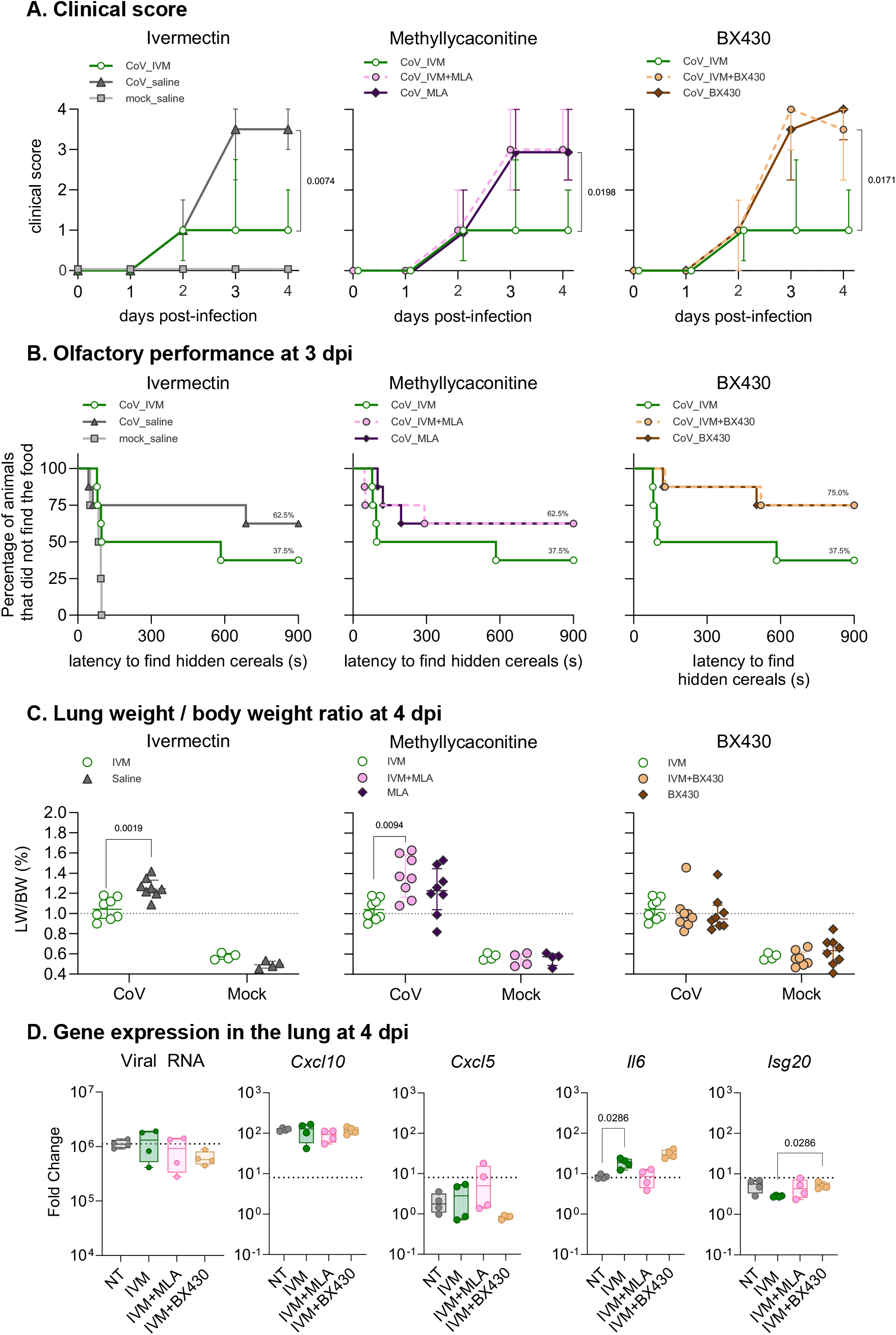
α7nAChR and P2X4 receptors mediate IVM-induced protection against SARS-CoV-2 in hamsters. Clinical profile of hamsters infected with SARS-CoV-2 and treated with nAChRs modulators and antagonists **(A)** Clinical score over four days post-infection (dpi) with GAT-107, EVP-6124, mecamylamine, methyllycaconitine or BX430 treatment. The clinical score is based on a cumulative 0-4 scale: ruffled fur; slow movements; apathy; and absence of exploration activity. **(B)** Olfactory dysfunction measured at 3 dpi of hamster either treated with GAT-107, EVP-6124, mecamylamine, methyllycaconitine or BX430. The olfaction test is based on the hidden (buried) food finding test. **(C)** Lung weight-to-body weight ratio measured at 4 dpi for hamsters treated with GAT-107, EVP-6124, mecamylamine, methyllycaconitine or BX430. (n=8 or 9/group). **(D)** Gene expression in lung. The expressions of viral RNA and *CXCL10*, *CXCL5, IL6*, and *ISG20* mRNA were measured. The values are normalized on non-infected non-treated conditions, only the infected conditions are shown. All the values are expressed as median, error bars indicate ± interquartile range. Mann-Whitney test was performed (P<0.05). The statistical test is performed between the non-treated condition and the different treated ones.

To functionally validate the contribution of α7nAChR, we combined IVM with either Mecamylamine (Meca), a non-competitive α7nAChR antagonist, or Methyllycaconitine (MLA), a competitive α7nAChR antagonist (Figure 5; Figure S3). Both Meca and MLA abolished the effect of IVM on clinical score (Figure 5A), olfactory dysfunction incidence (Figure 5B), and lung pathology (Figure 5C). Regarding P2X4 receptors, BX430 also abolished the beneficial effects of IVM on clinical scores and olfactory function (Figure 5A–B), but not on lung pathology (Figure 5C). The P2X7 antagonist could not be evaluated in this model owing to toxicity.

Altogether, these data demonstrate that the protective effect of IVM against SARS-CoV-2 in hamsters depends partially on both α7nAChR and P2X4 receptor signaling.

To bridge our *in vitro* and *in vivo* findings, we performed transcriptomic analysis of lung tissue from infected hamsters treated by the different drugs. Viral RNA levels in the lungs were comparable across all groups (Figure 5D), confirming that IVM treatment does not present an antiviral effect *in vivo*. No significant differences in *Cxcl10* or *Cxcl5* expression were observed between groups (Figure 5D). However, *Il-6* expression was significantly increased in IVM-treated animals compared with non-treated controls (p=0.0286, Mann-Whitney test), an effect that trend to decrease by co-treatment with MLA but not by BX430 (Figure 5D, S3). Finally, whereas IVM alone did not significantly alter *Isg20* expression, co-treatment with BX430 resulted in significant upregulation (Figure 5D, S3). Although the pulmonary transcriptional profile did not fully recapitulate the patterns observed in the *in vitro* co-culture model, the *in vivo* data further confirm that IVM-mediated protection against SARS-CoV-2–induced disease is dependent of both functional α7nAChR and P2X4 receptors.

## Discussion

Host-directed therapies remain strategically important for COVID-19 because they may retain activity across viral variants. The α7 nicotinic acetylcholine receptor (α7nAChR) emerged as a candidate target following epidemiological observations on smoking and disease severity, the so-called “nicotinic hypothesis” (37). Although vaccines have substantially reduced disease burden, therapeutic strategies specifically targeting the host inflammatory response remain limited. Given that severe COVID-19 is characterized by exaggerated pro-inflammatory cytokine production, modulating the host response represents a promising complementary approach. In parallel, purinergic signaling has emerged as another key regulator of inflammation (24).

IVM, a broad-spectrum anthelmintic, is also a positive allosteric modulator (PAM) of α7nAChR and purinergic receptors (16, 17, 19, 20). Clinical trials of IVM in COVID-19 have yielded conflicting results (43): whereas Chaccour et al. reported reduced anosmia when IVM was administered early (30), Reis et al. found no clinical benefit when treatment was initiated within seven days of symptom onset (32) indicating that there is no evidence of any positive effect of IVM in humans. However, these discrepancies and the previously demonstrated IVM improvement on the clinical outcome of hamsters infected with SARS-CoV-2 with a reduced IL-6/IL-10 ratio in the lungs (12) underscore the need to better characterize the molecular mechanisms underlying IVM’s effect.

Using two complementary *in vitro* systems: a monoculture of Vero E6 cells and a transwell co-culture model recapitulating epithelial–macrophage interactions in the lung, we highlighted that IVM protected cells from infection-induced death without significantly reducing viral RNA levels, indicating a host-directed rather than a direct antiviral mechanism. Transcriptomic analysis identified CXCL10 and CXCL5 as the most prominently regulated genes. In the transwell model, IVM induced rapid CXCL10 upregulation in THP-1 macrophages as early as 24 hours post infection (hpi), preceding epithelial overexpression at 48 hpi. CXCL10 is a key chemokine in antiviral defense, promoting the recruitment and activation of macrophages, lymphocytes, and NK cells prior to the establishment of adaptive immunity (44, 45). We therefore propose that early IVM-driven CXCL10 production in macrophages is a critical initiating event in the modulation of the host response.

Conversely, IVM markedly downregulated CXCL5 expression in both cell types. This finding appears at odds with a previous report showing that IVM increases CXCL5 secretion in primary human monocyte-derived macrophages via ATP-mediated activation of P2X4 receptors (46). One possible explanation is that, by preserving cell integrity, IVM limits the passive release of ATP into the extracellular space, thereby reducing P2X4 activation and subsequent CXCL5 induction. Alternatively, the difference may reflect distinct experimental systems (primary macrophages versus THP-1 cells) or timing of analysis. Nevertheless, our data suggest that P2X4 receptors contribute to the modulation of the inflammatory milieu during IVM treatment. Furthermore, data point to P2X4, rather than to α7nAChR or P2X7, as the primary mediator of IVM’s early transcriptomic effects *in vitro*. P2X4 and P2X7 receptors are known to form homo- and heterotrimers, with structural and functional crosstalk between them (25, 47, 48). Co-expression of P2X4 and P2X7 amplifies pro-inflammatory signaling and inhibits autophagy (26, 27), whereas P2X4 inactivation attenuates ATP-induced cell death (26). There is also evidence for functional coupling between α7nAChR and P2X7 through pannexin-1 (PANX-1) channels (34, 35, 49). It is therefore plausible that IVM modulates IL-6 and CXCL5 expression indirectly by altering the dynamics of this receptor network, although the exact topology of these interactions remains to be elucidated.

IVM is a type I PAM of α7nAChR, meaning that it potentiates receptor responses without altering desensitization kinetics (50). To compare different modes of α7nAChR activation, we selected two additional modulators from the ACRALL library: GAT-107, an ago-PAM, and EVP-6124, a competitive orthosteric agonist (39–42). Consistent with our previous report (12), IVM markedly improved clinical scores, reduced lung pathology, and alleviated anosmia. However, *in vivo*, not all α7nAChR modulators were equally effective: whereas the type I PAM (IVM) and the ago-PAM (GAT) improved outcomes, the competitive agonist EVP failed to ameliorate clinical signs. This difference likely reflects the distinct pharmacological profiles of these ligands. Orthosteric agonists such as EVP have been observed to induce a rapid and profound desensitized state in α7nAChR (51, 52), which may potentially limit the sustained signaling required for protection. In contrast, as a type I PAM, IVM does not activate α7nAChR by itself and therefore does not impose agonist-driven desensitization under sustained exposure; instead, it potentiates responses to endogenous agonists such as choline, thereby preserving α7nAChR signaling throughout treatment (50). The efficacy of GAT-107 aligns with this interpretation, as allosteric agonism by this ligand class is associated with slower activation kinetics and reduced desensitization compared with orthosteric activation (39, 41, 42, 52). Importantly, the protective effect of IVM was abolished when co-administered with either MLA or mecamylamine (Meca), confirming that functional α7nAChR signaling is necessary for IVM efficacy *in vivo*. Similarly, co-treatment with BX430 reversed IVM-induced clinical improvement, directly implicating P2X4 in the *in vivo* protective mechanism. Unfortunately, A438079 could not be tested in hamsters due to toxicity.

Strikingly, the *in vitro* and *in vivo* data diverged with respect to the role of α7nAChR. Whereas MLA did not reverse IVM’s transcriptomic signature in the transwell model, it completely abrogated protection in hamsters. Likewise, IL-6 expression is downregulated in THP-1 cells *in vitro* (as already described by (53)), whereas it is increased in the lungs of IVM-treated hamsters, as observed in patients with COVID-19 (6). These discrepancies are likely attributable to the greater cellular complexity of lung tissue *in vivo*, where additional cell types (including endothelial cells, fibroblasts, neurons, and dendritic cells) engage in crosstalk that is absent in our simplified two-cell transwell system. Under such conditions, the contribution of α7nAChR to the cholinergic anti-inflammatory pathway may become dominant, whereas the *in vitro* model highlights the leading role of P2X4 in macrophage-centered responses.

Collectively, our data indicates that IVM-mediated protection against SARS-CoV-2 pathology involves a dynamic and temporally coordinated interplay between cholinergic and purinergic signaling. Rather than acting through a single linear mechanism, IVM appears to orchestrate a multi-layered response in which early purinergic signaling initiates local immune recruitment, while later cholinergic signaling contributes to systemic resolution and tissue homeostasis. The most consistently regulated transcripts across models were CXCL5 and CXCL10, whose modulation may represent a central node through which IVM fine-tunes the balance between antiviral defense and immunopathology. We propose the following sequence of events (Figure 6): during the early phase of infection, IVM acts through P2X4 to induce rapid CXCL10 expression in macrophages. This early macrophage-derived signal likely serves to establish an initial antiviral chemotactic gradient, recruiting effector cells such as T lymphocytes and NK cells to sites of viral replication (54). At later time points, the primary source of CXCL10 appears to shift to epithelial cells, which may sustain or amplify this recruitment as the mucosal surface becomes the major site of pathogen recognition and containment. This transition from innate immune to epithelial-driven chemokine production suggests that IVM helps maintain a durable but controlled antiviral inflammatory environment rather than provoking an abrupt and potentially damaging cytokine burst. In parallel, by preserving epithelial cell viability and limiting virus-induced cytotoxicity, IVM may act upstream of P2X4 activation, limiting the lytic release of intracellular ATP and thereby reducing the extracellular ATP pool available for purinergic signaling. Because extracellular ATP is a potent danger-associated molecular pattern and P2X receptor agonist, its accumulation can fuel neutrophilic recruitment and pro-inflammatory signaling. By attenuating this signal, IVM concurrently dampens expression of CXCL5, a key neutrophil chemoattractant, and IL-6, a pleiotropic mediator of systemic inflammation. The net effect is a constrained inflammatory response in which antiviral recruitment via CXCL10 is preserved while excessive neutrophil influx and cytokine release. Importantly, the relative contribution of these pathways appears to differ between simplified *in vitro* systems and the more complex *in vivo* environment. While P2X4-dependent mechanisms are sufficient to explain many of our observations in cell-based models, functional α7nAChR signaling becomes indispensable for clinical protection in infected animals. This suggests that *in vivo*, IVM’s protective effects are integrated into broader neuro-immune regulatory circuits, such as the cholinergic anti-inflammatory pathway.

**Figure 6:**
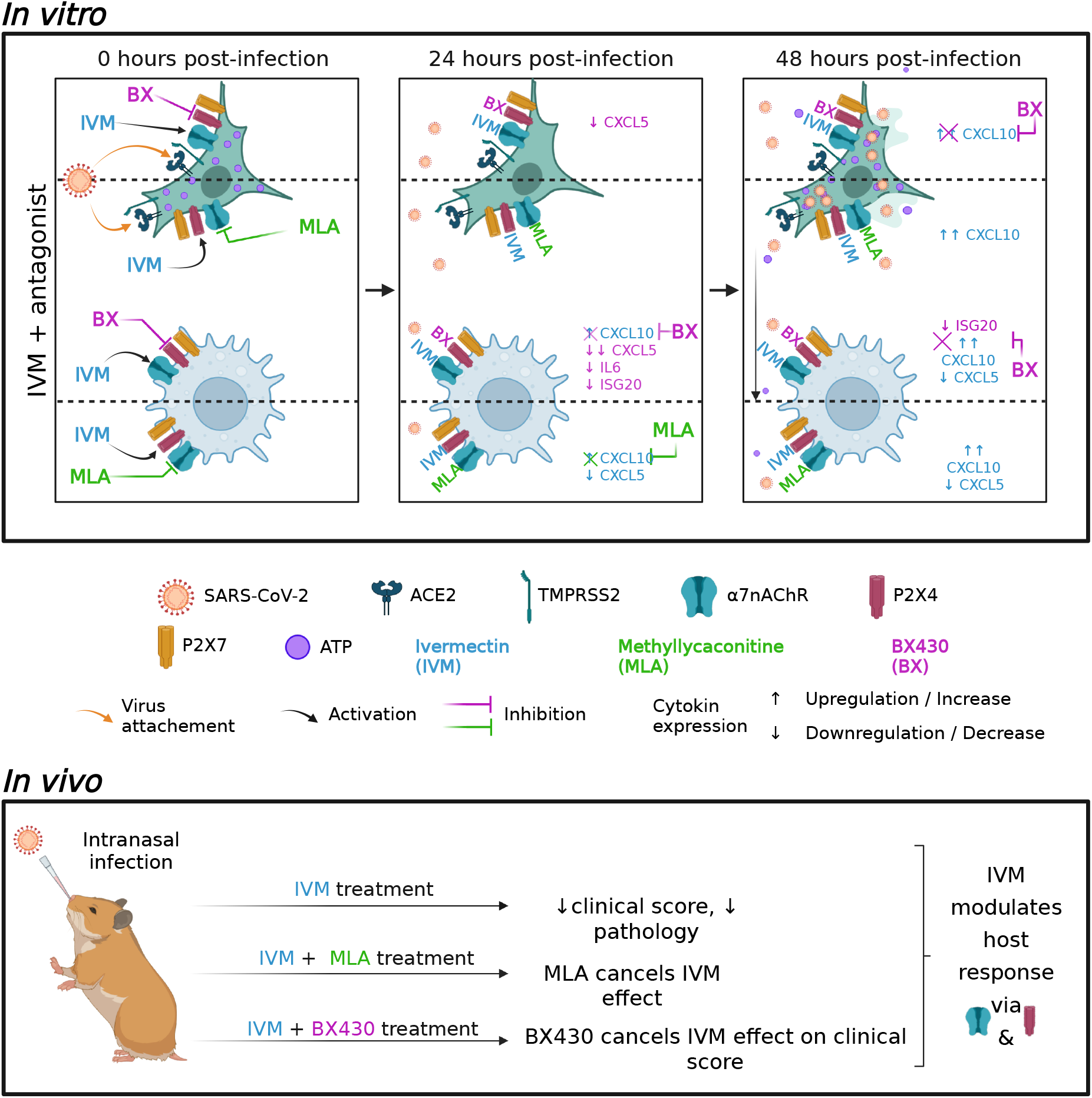
Overview of the *in vitro* and *in vivo* responses to purinergic and cholinergic modulators observed during SARS-CoV-2 infection. ***In vitro***, during infection, IVM induced through P2X4, an early and significantly higher expression of *CXCL10* by THP-1 cells, which is then highly produced by A549-ACE2-TMPRSS2 cells at 48 hpi. IVM induced an increase in cell survival and reduced the extracellular ATP concentration, consequently P2X4 is not ATP-activated, which could modify the profile of *IL6* and *CXCL5* mRNA expression and contain the antiviral response. ***In vivo***, IVM acts on both cholinergic receptors α7nACh and purinergic receptors to counteract the effect of SARS-CoV-2 infection.

In conclusion, our findings identify α7nAChR and P2X4 as functionally critical targets of IVM which through their sequential modulation in different type of cells (epithelial cells, macrophages and others as suggested by the *in vivo* analysis) induce a protective effect during SARS-CoV-2 infection. The convergence of cholinergic and purinergic pathways on the regulation of CXCL10 and CXCL5 highlights these chemokines as potential biomarkers and therapeutic targets in COVID-19. Given the central role of macrophages in coordinating the early inflammatory response, therapeutic strategies aimed at modulating these receptor networks (particularly through selective PAMs rather than orthosteric agonists) warrant further investigation.

## Materiel and methods

### Cells

The Vero E6 cells (ATCC #CRL-1586) were grown in Dulbecco’s Modified Eagle Medium (DMEM, #31966021, Gibco^TM^) medium supplemented with 5% fetal bovine serum. These cells were used for viral stock production, titration, and infection experiments.

The A549-ACE2-TMPRSS2 cells (supplied by the Virus and Immunity Unit hosted by Institut Pasteur and headed by Pr. Olivier Schwartz) were grown in Ham’s F-12K (Kaign) medium (#21127022, Gibco^TM^) supplemented with 10% fetal bovine serum and 1 µg/mL of blasticidine (#A1113903, Gibco^TM^).

The THP1 cells (provided by Dr G. Barba-Spaeth, Institut Pasteur) were cultivated in RPMI 1640 medium (Roswell Park Memorial Institute, #21875091, Gibco^TM^) supplemented with 10% fetal bovine serum. These cells, like the A549-ACE2-TMPRSS2 cells, were used in a co-culture system to study the role of α7AChRs and P2X receptors.

Cells were cultured in a humidified incubator at 37°C with 5% CO_2_. All cell lines were passed twice a week.

### SARS-CoV-2 virus and variants

The isolate BetaCoV/France/IDF00372/2020 (EVAg collection, Ref-SKU: 014V-03890) was kindly provided by Sylvie van der Werf. Alpha/B.1.1.7/IDF-IPP00078/2021 (GISAID ID: EPI_ISL_735391), Beta/B.1.351 (GISAID ID: EPI_ISL_964916) and Delta/B1.617.2 (GISAID ID: EPI_ISL_2029113) variants were supplied by the Virus and Immunity Unit hosted by Institut Pasteur and headed by Olivier Schwartz (Planas et al., 2021a, 2021b). Viral stocks were produced on Vero E6 cells infected at a multiplicity of infection of 1×10^-4^ PFU/cell. The virus was harvested 3 days post-infection, clarified and then aliquoted before storage at −80°C. Viral stocks were titrated on Vero E6 cells by classical plaque assays using semisolid overlays (Avicel, RC581-NFDR080I, DuPont) (55).

Wuhan_nLuc recombinant virus expressing Nanoluciferase (nLuc) as a reporter gene was obtained by reverse genetics as previously described (56).

### Modulators of cholinergic and purinergic receptors

Ivermectin (I8898, Sigma-Aldrich), a PAM for α7nAChR, P2X4 and P2X7, was used *in vivo* (one single subcutaneous injection) at the classical anti-parasitic dose of 400 µg/kg (Beco et al., 2001) simultaneously with infection and *in vitr*o at a dose of 0.625 µM, 1.25 µM, 2.25 µM and 5 µM. Mecamylamine (M9020, Sigma-Aldrich), a non-competitive α7nAChR antagonist, was used *in vivo* (one single intraperitoneal injection) at a dose of 1 mg/kg and *in vitr*o at a dose of 8 µM, 16 µM and 32 µM. Methyllycaconitine (M168, Sigma-Aldrich), a competitive α7nAChR antagonist, was used *in vivo* (one single intraperitoneal injection) at a dose of 1 mg/kg and *in vitr*o at a dose of 0.1 µM, 1 µM and 10 µM. GAT-107 (2621, Axon Medchem), an ago-PAM of α7nAChR, was used *in vivo* (one single intraperitoneal injection) at a dose of 0.08 mg/kg, 0.4 mg/kg, 2 mg/kg, 10 mg/kg and *in vitr*o at a dose of 6.25 µM, 12.5 µM and 25 µM. Encenicline, called EVP-6124 (HY-15430A/CS, MedChemExpress), an agonist of α7nAChR, was used *in vivo* (one single intraperitoneal injection) at a dose of 400 µg/kg and *in vitr*o at a dose of 2 µM, 4 µM, 8 µM, 16 µM. BX430 (SML1701, Sigma-Aldrich), an antagonist of P2X4, was used in *vitr*o at a dose of 5µM. A438079 (A9736, Sigma-Aldrich), an antagonist of P2X7, was used in vitro at a dose of 10µM.

### SARS-CoV-2 inhibition assay using Opera Phenix High-Content Screening System

Vero E6 cells were seeded in 96-well plates (655090, Greiner) in DMEM + 5% SVF and incubated overnight at 37°C and 5% CO_2_. 50µL/well of virus were inoculated for 1h at a multiplicity of infection (MOI) of 0.1. Then, the inoculum was removed and replaced by medium containing various concentrations of α7nAChRs modulators, prepared the same day. After 56 h or 72 h incubation at 37°C and 5% CO2, the plates were fixed with 4% PFA for 30 min at room temperature, then washed with PBS. The cells were then permeabilised for 10 min with 0.5% Triton X-100 and washed with PBS. To saturate non-specific sites, cells were incubated for 30 min with PBS + 5% Normal Goat Serum (#10000C, Thermofisher). The SARS-CoV-2 nucleoprotein was labelled overnight at 4°C with rabbit anti-SARS-CoV-2 Nucleocapsid primary antibody (#GTX135356, Clinisciences). After PBS washing, nuclei were labelled with Hoeschst 33342 (#62249, Thermofisher) for 15 min. Following a final wash, data was acquired using the OPERA Phenix High-Content Screening System to determine the number of cells per well and the percentage of infected cells.

### Co-culture *in vitro* model using Transwell

THP1 cells were seeded at the bottom of a Transwell plate (#3401, Costar) in RPMI medium + 10% SVF, in the presence of 100ng/mL PMA (P1585, Sigma-Aldrich), to differentiate them into macrophages. A549-ACE2-TMPRSS2 cells were seeded in the Transwell inserts, in FK12 Nut Mix + 10% SVF medium. After 24 h incubation, the medium was removed from the wells and inserts. 900 µL of RPMI medium + 10% SVF was added to THP1 cells and 450µL to uninfected A549-ACE2-TMPRSS2 cells; 250 µL of nLuc inoculate was added at a MOI of 0.1. After 1h incubation at 37°C and 5% CO2, the inoculate was removed and 450 µL of RPMI medium + 10% SVF was added to the inserts containing A549-ACE2-TMPRSS2 cells. Finally, RPMI medium containing various concentrations of α7nAChRs modulators or P2X receptor antagonists, prepared the same day, was added to the wells containing THP1 cells (100 µL) and to the inserts containing A549-ACE2-TMPRSS2 cells (50 µL). Infected cells were cultured for 24, 48 or 72 h depending of the experiments.

### Intracellular ATP measurements

In the previous transwell culture, the intracellular ATP was measured at 72 hours post-infection (hpi), in A549-ACE2-TMPRSS2 cells. The cells were lysed following the protocol of the ATPlite kit (6016941, PerkinElmer), ATP luminescence was measured in white-bottomed 96-well plates (655083, Greiner), using the VICTOR Nivo plate reader (Perkin Elmer).

### Quantification of the infection in the transwell cultures

The infection by Wuhan_nLuc was quantified by measuring nanoluciferase (nLuc) excreted in the culture medium for each part of the transwell. Twenty microliters of supernatant were mixed with same volume of nLuc subrate as indicated by the manufacturer (#N1110, Promega). nLuc measurement was performed in white-bottomed 96-well plates (655083, Greiner) using the VICTOR Nivo plate reader (Perkin Elmer).

### Transcriptomic analysis in transwell cultures

Transcriptomic analysis was performed on THP-1 cells only after 72 h of infection and on both THP-1 and A549-ACE2-TMPRSS2 after 24 and 48 hpi. The cells were lysed using RLT buffer (74104, Qiagen) at 72 hpi and the two types of cells were also lysed at 24 hpi and 48 hpi. RNA extraction was carried out using RNA esay kit (74104, Qiagen).

Samples were submitted to RT-qPCR. Briefly, RNA was reverse transcribed to first strand cDNA using the SuperScript™ IV VILO™ Master Mix (11766050, Invitrogen). qPCR was performed in a final volume of 10 µL per reaction in 384-well PCR plates using a thermocycler (QuantStudio 6 Flex, Applied Biosystems). Briefly, 2.5 µL of cDNA (12.5 ng) were added to 7.5 µL of a master mix containing 5 µL of Taqman (11732-020, Applied Biosystems) and 2.5 µL of nuclease-free water containing primer pairs (Table S1). The amplification conditions were as followed: 95°C for 10 min, 45 cycles of 95°C for 15 s and 60°C for 1 min; followed by a melt curve, from 60 °C to 95 °C. The *GAPDH* and the *18S* genes were used as reference. Variations in gene expression were calculated as the n-fold change in expression in the cells from the infected cells compared with the non-infected cellsgroup using the 2^-ΔΔCt^ method (57).

### Ethics

All animal experiments were performed according to the French legislation and in compliance with the European Communities Council Directives (2010/63/UE, French Law 2013–118, February 6, 2013) and according to the regulations of Institut Pasteur Animal Care Committees. This study (200023; APAFIS#25326-2020050617114340 v2) was approved by the Animal Experimentation Ethics Committee (CETEA 89) of the Institut Pasteur before experiments were initiated. Hamsters were housed by groups of four animals in isolators and manipulated in class III safety cabinets in the Pasteur Institute animal facilities accredited by the French Ministry of Agriculture for performing experiments on live rodents. All animals were handled in strict accordance with good animal practice.

### SARS-CoV-2 model in golden Syrian hamsters

Male golden Syrian hamsters (*Mesocricetus auratus;* RjHan:AURA) of 5-6 weeks of age (average weight 60-80 grams) were purchased from Janvier Laboratories and handled under specific pathogen-free conditions. Before any manipulation, animals underwent an acclimation period of one week. The animals were housed and manipulated in isolators in a biosafety level-3 facility, with *ad libitum* access to water and food. Animals were anesthetized with an intraperitoneal injection of 200 mg/kg ketamine (Imalgène 1000, Merial) and 10 mg/kg xylazine (Rompun, Bayer). 100 µL of physiological solution containing 6×10^4^ PFU of SARS-CoV-2 (Wuhan) was administered intranasally to each animal (50 µL/nostril). Mock-infected animals received the physiological solution only. Infected and mock-infected hamsters were housed in separate isolators and were followed-up daily during four days at which the body weight and the clinical score were noted. The clinical score was based on a cumulative 0-4 scale: ruffled fur, slow movements, apathy, absence of exploration activity. At day 3 post-infection (dpi), animals underwent a food finding test to assess olfaction as previously described (12, 58). Briefly, 24 hours before testing, hamsters were fasted and then individually placed into a fresh cage (37 x 29 x 18 cm) with clean standard bedding for 10 minutes. Subsequently, hamsters were placed in another similar cage for 2 minutes when about 5 pieces of cereals were hidden in 1.5 cm bedding in a corner of the test cage. The tested hamsters were then placed in the opposite corner and the latency to find the food (defined as the time to locate cereals and start digging) was recorded using a chronometer. The test was carried out during a 15 min period. As soon as food was uncovered, hamsters were removed from the cage. The tests were carried out in isolators in a Biosafety level-3 facility that were specially equipped for that. At 4 dpi, animals were euthanized with an excess of anesthetics (ketamine and xylazine) and exsanguination (12).

### RNA isolation from golden hamsters’ lungs

Lungs from infected and treated animals were recovered at 4 dpi and frozen in several fragments. Frozen lung fragments were weighted and homogenized with 1 mL of ice-cold DMEM supplemented with 1% penicillin/streptomycin (15140148, Thermo Fisher) in Lysing Matrix M 2 mL tubes (116923050-CF, MP Biomedicals) using the FastPrep-24™ system (MP Biomedicals), and the following scheme: homogenization at 4.0 m/s during 20 sec, incubation at 4 °C during 2 min, and new homogenization at 4.0 m/s during 20 sec. The tubes were centrifuged at 10,000 × *g* during 2 min at 4 °C, and the supernatants collected. 125 µL of the supernatants were homogenized with 375 µL of Trizol LS (10296028, Invitrogen).

### Transcriptomics analysis in golden hamsters’ tissues

Total RNA was extracted from the lungs using the Direct-zol RNA MiniPrep kit (R2052, Zymo Research). 125 µL of tissue homogenate was incubated with 375 µL of Trizol LS (10296028, Invitrogen) and the extraction was performed according to the manufacturer’s instructions.

RNA preparations from lungs collected at 4 dpi were submitted to RT-qPCR. Briefly, RNA was reverse transcribed to first strand cDNA using the SuperScript™ IV VILO™ Master Mix (11766050, Invitrogen). qPCR was performed in a final volume of 10 µL per reaction in 384-well PCR plates using a thermocycler (QuantStudio 6 Flex, Applied Biosystems) and its related software (QuantStudio Real-time PCR System, v.2.7.0, Applied Biosystems). In each well, 2.5 µL of cDNA (12.5 ng) was added to 5 µL of Power SYBR Green PCR Master Mix (4367659, Applied Biosystems) and 2.5 µL of nuclease-free water containing 1 µM of golden hamster primer pairs (Supplementary Table S1;(4, 56)). The amplification conditions were as follows: 50°C for 2 min, 95 °C for 10 min, and 45 cycles of 95°C for 15 s and 60°C for 1 min, followed by a melt curve from 60°C to 95°C. The *γ-actin* and the *Hprt* (hypoxanthine phosphoribosyl-transferase) genes were used as reference. Variations in gene expression were calculated as the n-fold change in expression in the tissues from the infected hamsters compared with the tissues of the mock-infected group using the 2-ΔΔCt method (57).

### Statistics

Statistical tests were carried out using Prism software (GraphPad Prism, version 10, San Diego, USA), with p < 0.05 considered significant. Quantitative data was compared across groups using Kruskal-Wallis followed by the Dunn’s multiple comparisons test and Mann-Whitney. Randomization and blinding were not possible due to pre-defined housing conditions (separated isolators between infected and non-infected animals).

## Data availability

Data will be made available on request.

## Acknowledgements

The SARS-CoV-2, strain BetaCoV/France/IDF00372/2020 (EVAg collection, Ref-SKU: 014V-03890) strain was supplied by the National Reference Centre for Respiratory Viruses hosted by Institut Pasteur (Paris, France) and headed by Pr. Sylvie van der Werf. The human sample from which strain 2019-nCoV/IDF0372/2020 was isolated has been provided by Dr. X. Lescure and Pr. Y. Yazdanpanah from the Bichat Hospital (Paris, France). The isolate BetaCoV/France/IDF00372/2020 was kindly provided by Pr. Sylvie van der Werf. Alpha/B.1.1.7/IDF-IPP00078/2021, Beta/B.1.351 and Delta/B1.617.2 variants, and the A549-ACE2-TMPRSS2 cells were supplied by the Virus and Immunity Unit hosted by Institut Pasteur and headed by Pr. Olivier Schwartz. The ACRALL library was compiled by Federica Brando.

Part of this work was performed at the UtechS Photonic BioImaging (PBI) platform supported by Institut Pasteur and by Région Ile-de-France.

This work was supported by the Fondation pour la Recherche Médicale (grant ANRS MIE202112015304). V.P. is recipient of a fellowship from the European Union’s Horizon 2020 Framework Program for Research and Innovation under Specific Grant Agreement No. 945539 (Human Brain Project SGA3).

## Supplementary figures

**Figure S1:**
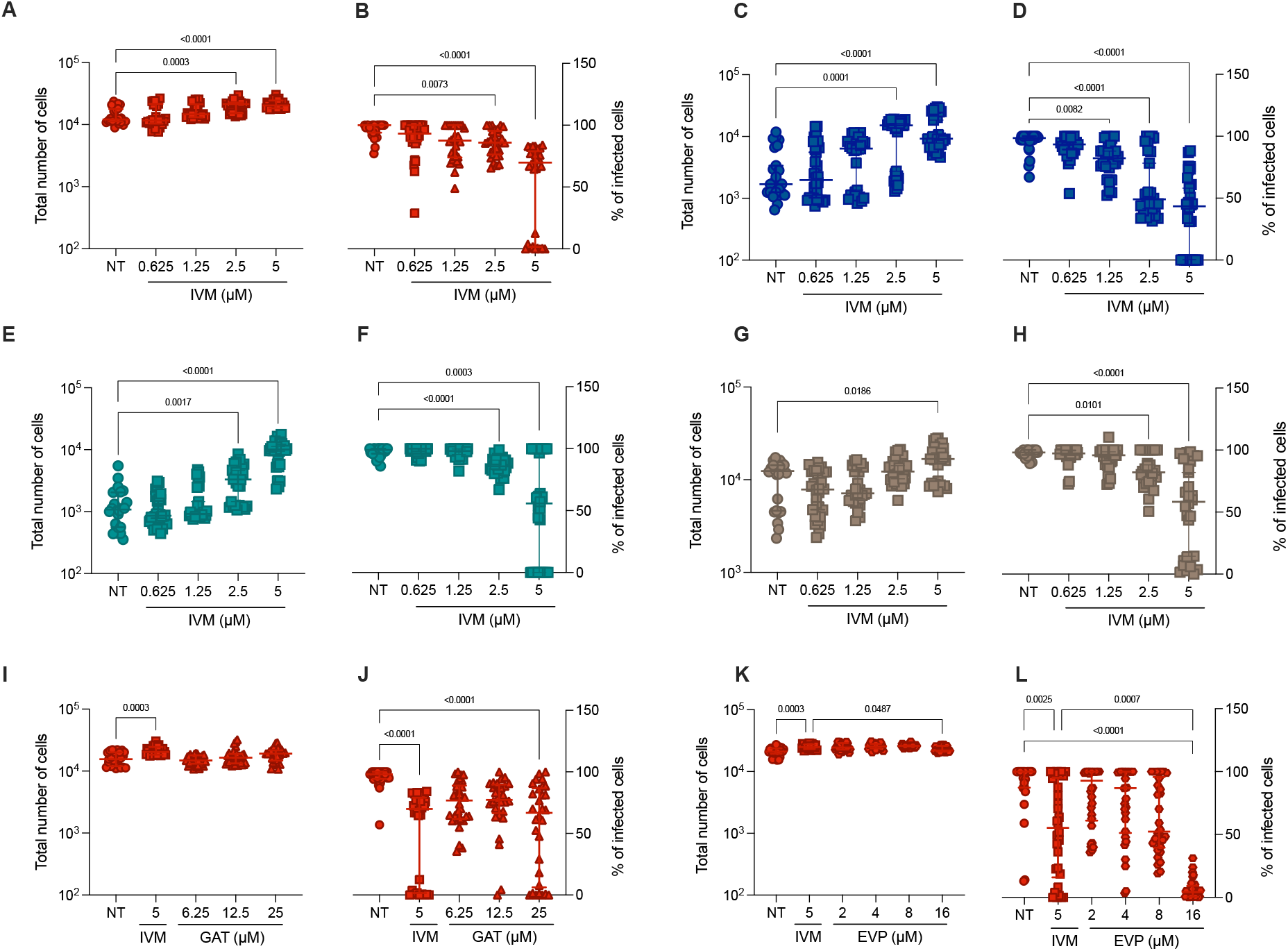
Cholinergic receptors modulators effect is variant independent in an *in vitro* model. In the Vero E6 *in vitro* model, with the Opera Phenix High Content Screening System technology, **(A, C, E, G, I, K)** total number of cells or **(B, D, F, H, J, L)** percentage of infected cells was measured in the presence of several α7nAChR modulators at various concentration: ivermectin (IVM, at 0.625 µM, 1.25 µM, 2.5µM and 5 µM), GAT-107 (GAT, at 6.25 µM, 12,5 µM and 25 µM) and EVP-6124 (EVP, at 2 µM, 4 µM, 8 µM and 16 µM). Vero E6 cells were infected at an MOI of 0.1 during 56 h for the Wuhan virus (in red) or 72 h for Alpha (in blue), Beta (in green) and Delta variants (in brown). These datas enabled us to determine the dose of the drug that had the greatest effect on cell survival and the percentage of infected cells. All the values are expressed as means, error bars indicate ± SD. Three independent experiments with 10 technical replicates were performed with the Opera Phenix technology. Kruskall-Wallis test followed by the Dunn’s multiple comparisons test (only the significant values are shown).

**Figure S2:**
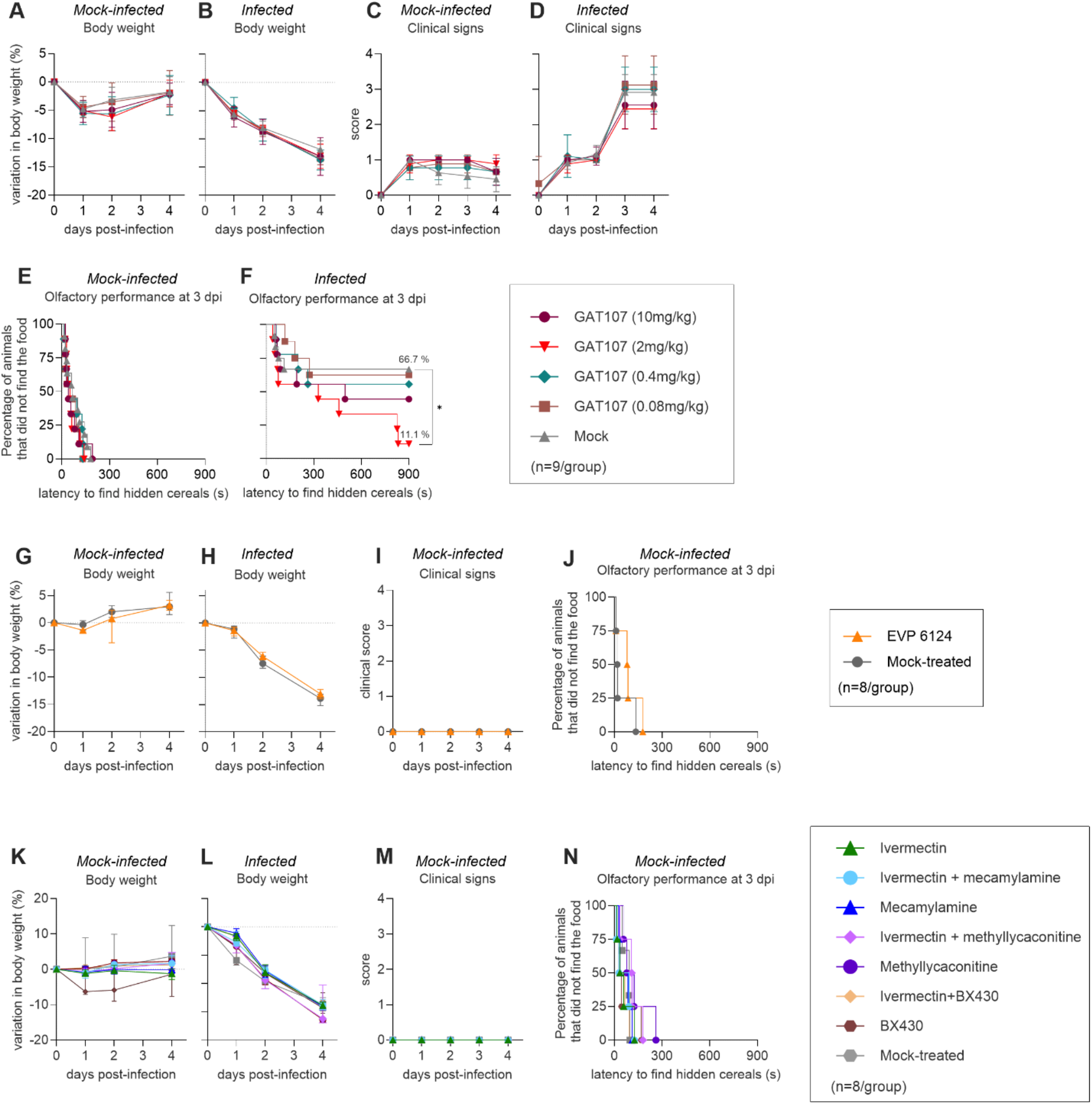
Clinical profile of hamsters infected with SARS-CoV-2 and treated with nAChRs agonists (GAT-107 or EVP-6124) or nAChRs antagonists (mecamylamine or methyllycaconitine). **(A, B, G, H, K, L)** Body weight variation over four days post-infection (dpi) in **(A, G, K)** mock-infected and **(B, H, L)** infected hamsters. **(C, D, I, M)** Clinical score over four days post-infection of: **(C, I, M)** mock-infected hamsters or **(D)** infected hamsters treated with different doses of GAT-107 (10 mg/kg; 2 mg/kg; 0.4 mg/kg; 0.08 mg/kg). The clinical score is based on a cumulative 0– 4 scale: ruffled fur; slow movements; apathy; and absence of exploration activity. **(E, F, J, N)** Olfactory performance measured at 3 dpi. The olfaction test is based on the hidden (buried) food finding test. Curves represent the olfactory performance of animals during the test either mock-infected **(E, J, N)** or **(F)** infected treated with with mecamylamine, methyllycaconitine, GAT-107 or EVP-6124. Horizontal lines indicate median and the interquartile range. Kruskal-Wallis test followed by the Dunn’s multiple comparisons test was performed (*P<0.05, **P<0.01, ***P<0.001, ****P<0.0001). (n=8 or 9/group).

**Figure S3:**
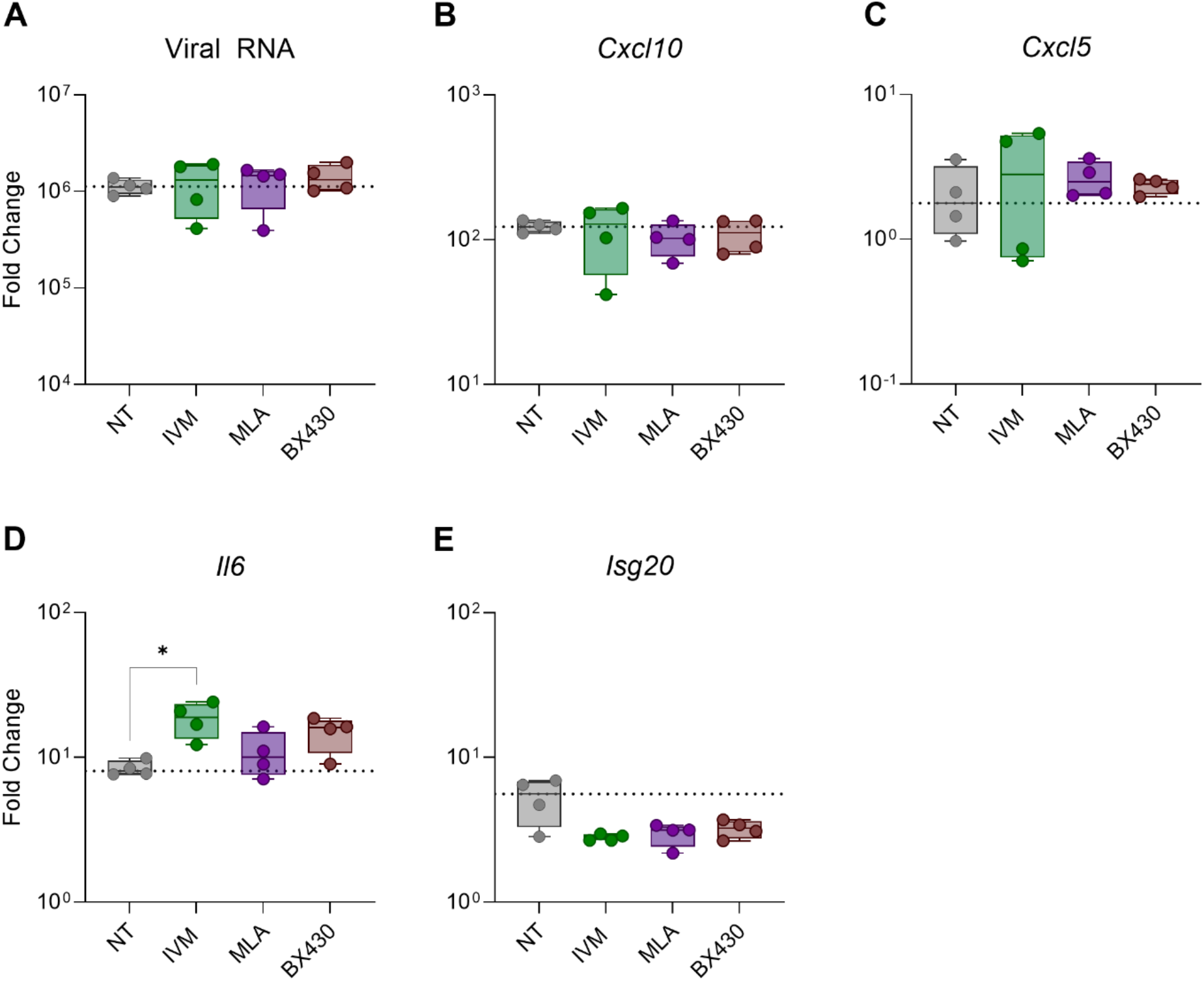
*In vivo* modulation of purinergic or α7-nAChRs receptors modulates hamsters host-immune response to SARS-CoV-2. Gene expression values in the lung of SARS-CoV-2 infected hamsters treated with IVM or with cholinergic and purinergic antagonists: BX430 (at 1 mg/kg, antagonizing P2X4), and methyllycaconitine (MLA, at 1 mg/kg, antagonizing α7nAChR). The mRNA expression of **(A)** Viral RNA, **(B)** *Cxcl10*, **(C)** *Cxcl5*, **(D)** *Il6*, and **(E)** *Isg20* were measured. The values are normalized on infected non-treated conditions, only the infected conditions are shown. Four independent experiments were performed in technical duplicates. All the values are expressed as median, error bars indicate ± interquartile range. Mann-Whitney test was performed (*P<0.05). The statistical test is performed between the non-treated condition and the different treated ones.

## Supplementary Table

**Supplementary Table 1:**
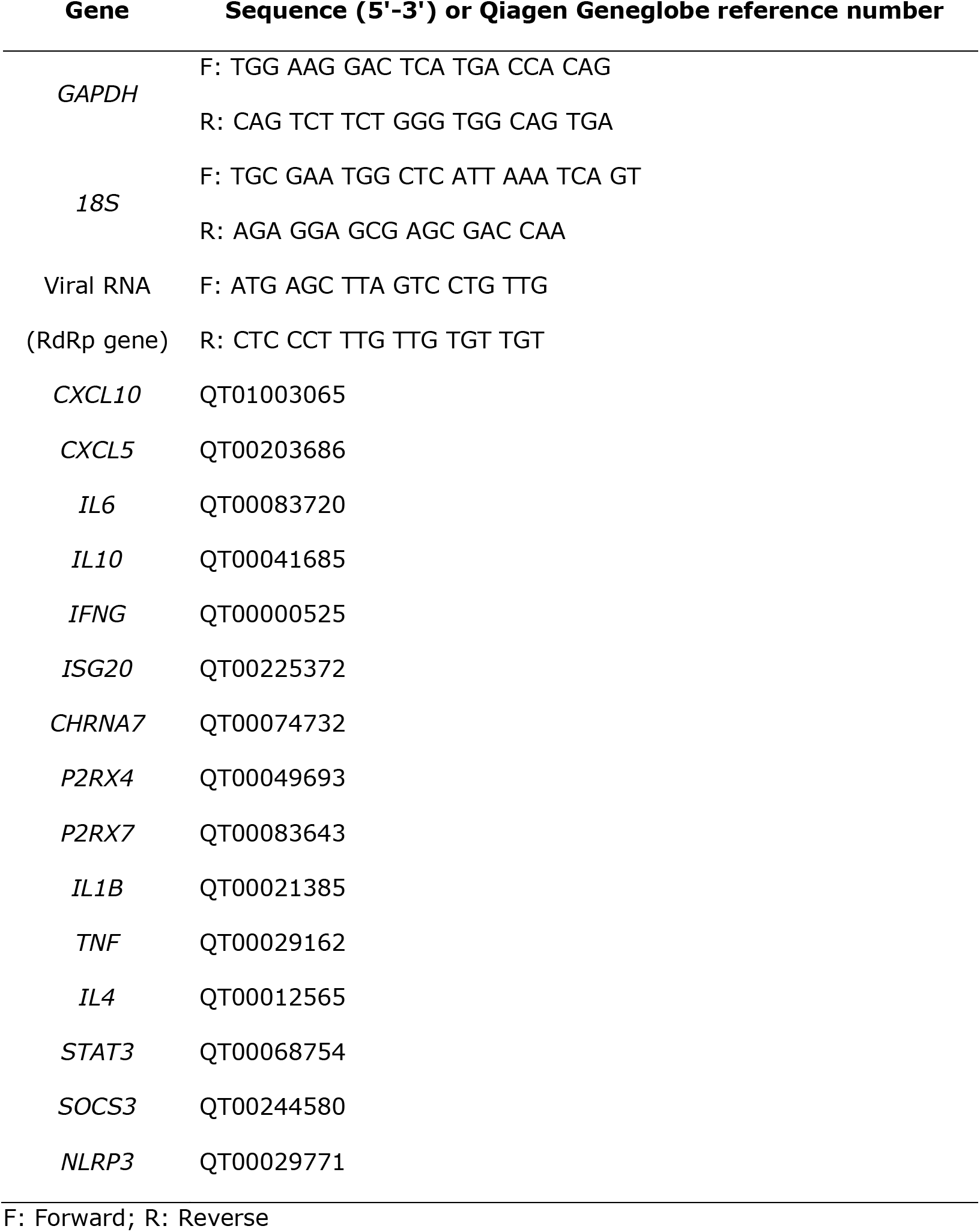
List of primers used in the transcriptomic analysis.

